# Biomarkers of collagen synthesis predict progression in the PROFILE idiopathic pulmonary fibrosis cohort

**DOI:** 10.1101/497156

**Authors:** Louise A Organ, Anne-Marie R Duggan, Eunice Oballa, Sarah C Taggart, Juliet K Simpson, Arthur R Kang’ombe, Rebecca Braybrooke, Philip L Molyneaux, Diana J Leeming, Morten A Karsdal, Carmel B Nanthakumar, William A Fahy, Richard P Marshall, R Gisli Jenkins, Toby M Maher

## Abstract

Idiopathic pulmonary fibrosis (IPF) is characterised by excessive extracellular matrix (ECM) deposition and remodelling. Measuring this activity provides an opportunity to develop tools capable of identifying individuals at-risk of progression. Longitudinal change in markers of ECM synthesis was assessed in 145 newly-diagnosed individuals with IPF.

Serum levels of collagen synthesis neoepitopes, PRO-C3 and PRO-C6 (collagen type 3 and 6), were elevated in IPF compared with controls at baseline, and progressive disease versus stable disease during follow up, (PRO-C3 p<0.001; PRO-C6 p=0.029). Assessment of rate of change in neoepitope levels from baseline to 3 months (defined as ‘slope to month 3’: HIGH slope, slope > 0 vs. LOW slope, slope <=0) demonstrated no relationship with mortality for these markers (PRO-C3 (HR 1.62, p=0.080); PINP (HR 0.76, p=0.309); PRO-C6 (HR 1.14, p=0.628)). As previously reported, rising concentrations of collagen degradation markers C1M, C3M, C6M and CRPM were associated with an increased risk of overall mortality (HR=1.84, CI 1.03 – 3.27, p=0.038, HR=2.44, CI 1.39–4.31, p=0.002; HR= 2.19, CI 1.25–3.82, p=0.006; HR= 2.13 CI 1.21–3.75, p=0.009 respectively).

Elevated levels of PRO-C3 and PRO-C6 associate with IPF disease progression. Collagen synthesis and degradation biomarkers have the potential to enhance clinical trials in IPF and may inform prognostic assessment and therapeutic decision making in the clinic.

## Introduction

Idiopathic pulmonary fibrosis (IPF) is a fatal condition with a dismal untreated prognosis and despite the approval of two anti-fibrotic therapies remains a major clinical challenge. At present there are no available measures for stratifying individuals with IPF to specific therapies nor, importantly, to determine response to therapy. Furthermore, although forced vital capacity (FVC) is accepted as a regulatory end-point in clinical trials, the relative insensitivity of physiological measures to change in fibrosis necessitates studies with a minimum duration of 52 weeks [1]. Therefore, there is an urgent need to identify biomarkers for various aspects of disease behaviour in IPF; particularly those that detect individuals at greater risk of progression and which stratify individuals into better defined endotypes (as a path to enabling effective targeting of therapies to individuals most likely to benefit).

Excessive extracellular matrix (ECM) deposition is central to the pathogenesis of IPF. The production, deposition and remodelling of collagen, a major component of ECM, is a dynamic process [2]. In healthy lung, there is a balance between degradation and synthesis of collagen and other ECM proteins [2], which is disturbed in the lungs of individuals with fibrosis [3–5]. During collagen turnover, distinct peptides are generated as by-products of both collagen breakdown and synthesis. The pro-peptide of pro-collagen is cleaved before collagen is incorporated into the matrix. This results in the release of unique fragments for each individual form of collagen [6]. Similarly, during degradation, specific metalloproteases (MMPs) cleave collagen fibres revealing distinct neo-epitopes. Both synthesis peptides and cleavage neo-epitopes are released into the circulation and can be detected in the blood.

We have previously demonstrated, in subjects recruited to the PROFILE (Prospective Observation of Fibrosis in the Lung Clinical Endpoints) study, that serum levels of neoepitopes are higher in individuals with IPF when compared with healthy age-matched controls. Furthermore, we demonstrated that longitudinal changes in serum concentrations of several neo-epitopes track with progression of fibrosis and, by three months, can predict mortality in individuals with IPF [7]. In the current study, we assessed collagen formation markers for their prognostic utility in IPF in a separate cohort of individuals recruited into the PROFILE study. We also compared these synthesis markers to their collagen-equivalent degradation neoepitopes thus enabling assessment of the relationship between synthesis and degradation during IPF progression.

## Methods

### Study design and participants

As previously described, the PROFILE study is a prospective, multicentre, observational cohort study that recruited individuals with IPF and fibrotic NSIP from two co-ordinating UK centres; Nottingham and Royal Brompton [7, 8]. Participants were enrolled if they had an incident diagnosis confirmed at multidisciplinary discussion according to international consensus guidelines [9]. The cohort of individuals used in the current study represents a sub-group of 145 participants from PROFILE; none of whom were included in the previous PROFILE study on neoepitopes [7] and all of whom had at least 1 year of follow up at the time neoepitope analyses were performed. A total of 50 physician-verified control participants without history of respiratory disease were recruited from primary care clinics. Samples from 20 of these subjects were used in our earlier study [7]. All participants provided written informed consent. The PROFILE study is registered on ClinicalTrials.gov, numbers NCT01134822 and NCT01110694. For survival analyses, a censoring date of 1^st^ March 2016 was used.

## Procedures

### Patient samples

Serum samples collected from study participants at baseline, 1, 3, and 6 months, and a single serum sample from controls were stored at −80 °C until assayed. Study-specific operating procedures for collection of the serum were used to ensure standardisation across both centres. Briefly, blood was collected with anticoagulant free, serum separation tubes, with coated silica as the clot activator (Becton Dickenson, Winnersh, UK). All samples were processed within 2h of collection. Samples were allowed to clot at RT for 30min and serum was then isolated through centrifugation, and aliquoted before freezing.

### Quantification of extracellular matrix related biochemical markers

Serum samples from patients were analysed using competitive ELISAs for the formation markers of type 1 collagen (P1NP) [10], type 3 collagen (PRO-C3) [11] and type 6 collagen (PRO-C6) [12] as previously described. Additionally, assessment of matrix metalloproteinase (MMP) degraded biglycan (BGM), - type 1 collagen (C1M) [13], - type 3 collagen (C3M) [14], - type 6 collagen (C6M) [15] and C-reactive protein - (CRPM) [16]; was undertaken as previously described [7]. A single aliquot for each subject at each time-point was used for all assays to avoid repeated freeze/thaw of samples. Samples were analysed blinded to the associated clinical data. Performance characteristics of the assays and non-standard abbreviations are described in the online data supplement, Table 1E.

**Table 1.** Baseline clinical characteristics in participants with idiopathic pulmonary fibrosis (n=145) and healthy controls (n=50) Data are mean (SD) or n (%), unless otherwise stated. BMI=body-mass index. FVC=forced vital capacity. DLCO=diffusion capacity for carbon monoxide. CPI=composite physiological index, *Number of subjects analysed due to missing data

|  | **IPF subjects (n=145)** |
| --- | --- |
| **Age (years)** | 71.7(7.7) |
| **Sex (male)** | 117 (81%) |
| **Ethnicity** |  |
| *Caucasian* | 142 (97.9%) |
| *Asian* | 2 (1.4%) |
| *Mixed Race* | 1 (0.7%) |
| **Ever smokers** | 108 (74%) |
| **BMI (kg/m<sup>2</sup>)** | 28.4 (4.4) *n=144 |
| **FVC (% predicted)** | 79.8 (20.4) *n=140 |
| **DLCO (% predicted)** | 48.2 (17.9) *n=129 |
| **CPI** | 44.9 (14.9) *n=129 |
| **Median follow- up days** | 1051 (422.9) |
|  | **Healthy controls (n=50)** |
| **Age (years)** | 64.6 (7.27) |
| **Sex (male)** | 40 (80%) |

### Lung function measurements

Lung function measurements for forced vital capacity (FVC) and diffusion capacity for carbon monoxide (DLCO) were performed on participants at baseline, 6 months and 12 months according to current ATS/ERS guidelines. Percent-predicted values are based on the European Coal and Steel Board reference equations using age and height at the baseline visit [17].

### Statistical Analysis

As previously described, ‘progressive disease’ was defined as ≥10% decline in FVC and/or death within 12 months. Missing lung function data were not imputed. Where no lung function data beyond baseline was available and the subject had not died within the relevant period, progression status was adjudicated by the responsible physician, after case note review and blinded to biomarker data at the time of adjudication. A total of 9 subjects were adjudicated for progression: 1 of these subjects were adjudicated as progression and 8 were adjudicated as stable.

In cases where biomarker data were below the lower limit of detection, values were imputed to be half the lower limit of detection. Values above the upper limit were conservatively imputed as the upper limit of detection. Imputation rates ranged between 0.9% to below 10% for all markers except C3M that had approximately 19% imputation. Before statistical analysis all biomarkers were transformed (log10 transformed where the biomarker is treated as response and log2 where the biomarker is treated as a covariate).

A linear regression model adjusted for gender and age was used to evaluate the association between patient group (healthy controls and IPF subjects) and biomarker values at baseline. Adjusted estimates of group means, 95% CIs and p-values of the estimated mean difference are reported. For serial biomarker data, change from baseline for each participant is represented by a gradient on a linear model where the single explanatory variable is the number of months between the baseline visit and subsequent visits using the recorded visit date. We used a linear mixed-effects model (Mixed Model Repeated Measures (MMRM)) to investigate associations between repeated continuous biomarker response and progression status (progression vs stable), visit, visit by progression status (controls are not included in the model). This analysis was adjusted for age, site, gender and smoking status. Unstructured covariance was assumed for all repeated biomarker response values within a participant. Adjusted estimates of biomarker means (95% CIs) by progression status at each visit and p-values of the test of mean differences between progression vs. stable at each visit are reported. Apart from biomarker data above or below the limits of quantification, all other missing biomarker data was not imputed and was assumed Missing at Random (MAR) implying that after adjusting for important covariates in the model the missing data was independent of the observed data.

Univariate analysis was conducted to examine the effect of baseline biomarker measures on overall survival. Two separate procedures were used; the Kaplan-Meier method for survival curves and a proportional hazards model to derive Hazard Ratios.

A proportional hazards model was used to evaluate association between time to event (overall survival) as response and continuous biomarker value as a covariate. Similarly, in a separate model, baseline % predicted FVC and separately % predicted DLco were included as covariates. A proportional hazards model was used to evaluate association between time to event (overall survival) as response and biomarker slope as a factor (defined as ‘slope to month 3’: HIGH (rising concentrations, slope > 0) versus LOW (stable or falling concentrations, slope <=0). Hazard ratios represent either risk of mortality for a 2-fold increase in the biomarker value or risk of mortality in participants with rising neoepitope concentrations relative to those with stable or falling concentrations.

Associations between explanatory variables and endpoints were deemed to be significant at the 5% level. All statistical analyses were conducted with SAS (version 9.3) and were independently quality controlled by a second statistician.

## Results

145 IPF participants recruited in the PROFILE study were assessed in the current analysis. As expected, participants were predominantly male (81%) with a mean age of 71.7 years (SD 7.7) and moderate lung function impairment; FVC 79.8 (SD 20.4) %predicted and DLco 48.2 (SD 17.9) %predicted. Subject demographics are presented in Table 1. The 50 control subjects were 80% male with a mean age of 64.6 (SD 7.27).

At baseline, levels of PRO-C3 and PRO-C6 but not P1NP, were significantly elevated in individuals with IPF compared with healthy controls (Figure 1). To enable paired assessment of collagen synthesis and degradation, levels of neoepitopes previously linked with disease progression and survival in the PROFILE study [7] were measured. These included the three degradation neoepitopes which pair with the synthesis markers assessed; C1M (pairs with P1NP), C3M (PRO-C3) and C6M (PRO-C6) and two additional neoepitopes from our previous study, CRPM and BGM. In keeping with our previously presented data, C1M, C3M, C6M and CRPM neoepitopes were significantly elevated in the serum of subjects with IPF compared with controls at baseline (Figure 1E).

**Figure 1.**
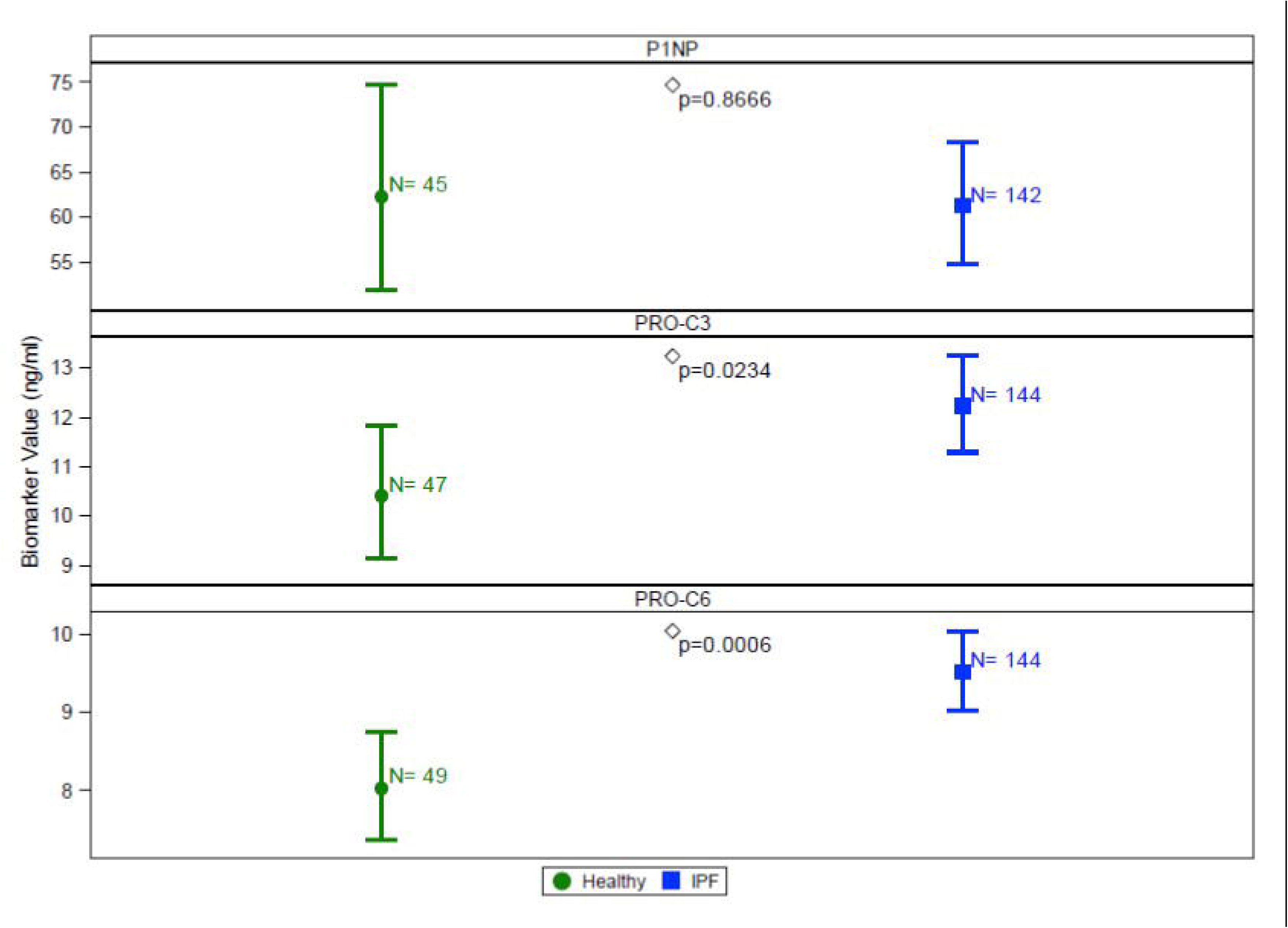
Baseline comparison of collagen synthesis neoepitope concentrations for collagen-type 1 (P1NP), - type 3 (PRO-C3) and - type 6 (PRO-C6) in healthy controls and participants with idiopathic pulmonary fibrosis. Plots represent mean and 95% CI (error bars) adjusted for age and gender.

73 subjects had progressive disease whilst 71 had stable disease. Levels of both PRO-C3 and PRO-C6 were higher in individuals with progressive disease compared with those with stable disease at baseline and they remained consistently and significantly higher in individuals with progressive disease at all visits (1,3 and 6 months) (Figure 2). There were no significant differences between stable and progressive disease at any visit for P1NP (Figure 2).

**Figure 2.**
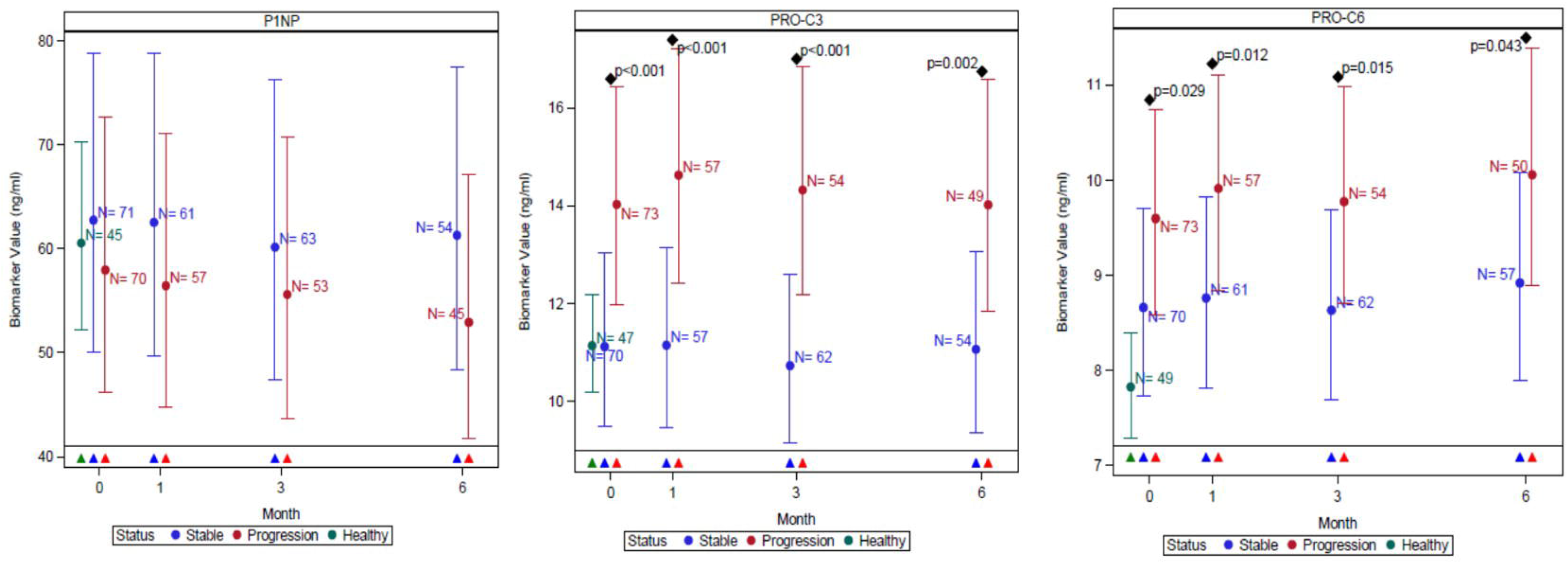
Comparisons of neoepitope concentrations for collagen synthesis markers in healthy control and participants with stable and progressive IPF, measured at baseline, 1, 3 and 6 months after baseline for collagen-type 1 (P1NP), - type 3 (PRO-C3) and - type 6 (PRO-C6). Plots represent mean and 95% CI (error bars) adjusted for age, sex, site and smoking status. Disease progression was defined as all-cause mortality or ≧10% decline in forced vital capacity at 12 months. The number of evaluable samples available for analysis at each time point are provided in the graph. P values are provided where significant (p<0.05) differences were observed between stable and progressive disease at a particular time point. Triangles represent which groups were compared at each time point on the graph, based on colour.

Consistent with our previously reported findings, C1M, C3M, C6M and CRPM were also significantly higher in those with progressive disease compared with individuals with stable disease across all time-points (baseline, 1, 3 and 6 months) (see Figure 2E in the online data supplement).

When comparing ratios of synthesis and degradation markers P1NP:C1M was significantly lower in participants with progressive disease from 1 month onwards compared with stable disease, whilst PRO-C3:C3M was significantly lower in participants with stable disease at each time point compared with progressive disease (Figure 3). There were no significant differences between stable and progressive disease at any timepoint for the ratio of PRO-C6:C6M (Figure 3).

**Figure 3.**
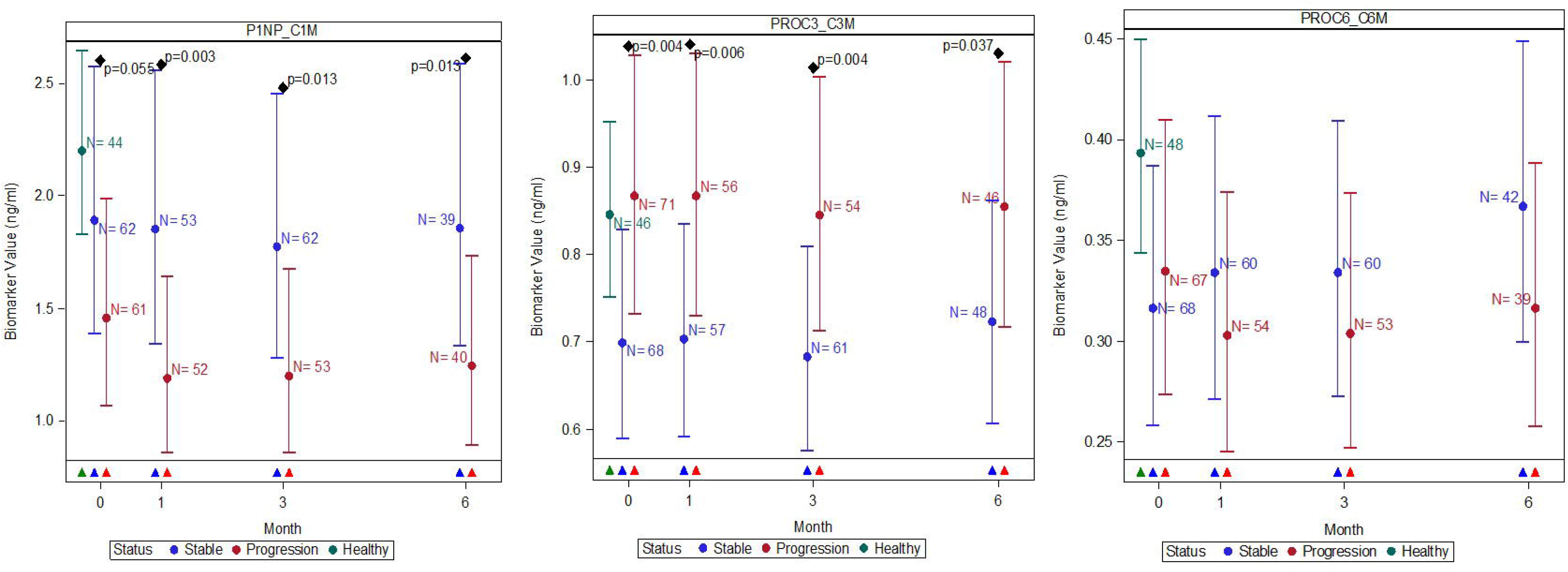
Comparisons of absolute values in the ratio of neoepitope collagen synthesis to degradation markers in healthy controls and participants with stable and progressive IPF, measured at baseline, 1, 3 and 6 months for: collagen-type 1(P1NP:C1M), - type 3 (PRO-C3:C3M) and - type 6 (PRO-C6:C6M). Plots represent mean and 95% CI (error bars) adjusted for age, sex, site and smoking status. Disease progression was defined as all-cause mortality or ≧10% decline in forced vital capacity at 12 months. The number of evaluable samples available for analysis at each time point are provided in the graph. P values are provided where significant (p<0.05) differences were observed between stable and progressive disease at a particular time point. Triangles represent which groups were compared at each time point on the graph, based on colour.

In an unadjusted univariate analysis, baseline concentrations of PRO-C3 (H.R. 1.49 (1.04-2.13), p = 0.030) and P1NP:C1M ratio (H.R. 0.67 (0.53-0.85), p=0.001) were predictive of survival (Table 2). As expected, % predicted FVC at baseline (H.R. 0.22 (0.12-0.39, p<0.0001) and % predicted DLco at baseline (H.R. 0.16 (0.09-0.30), p<0.0001) were strongly predictive of survival (Table 2.).

**Table 2.**
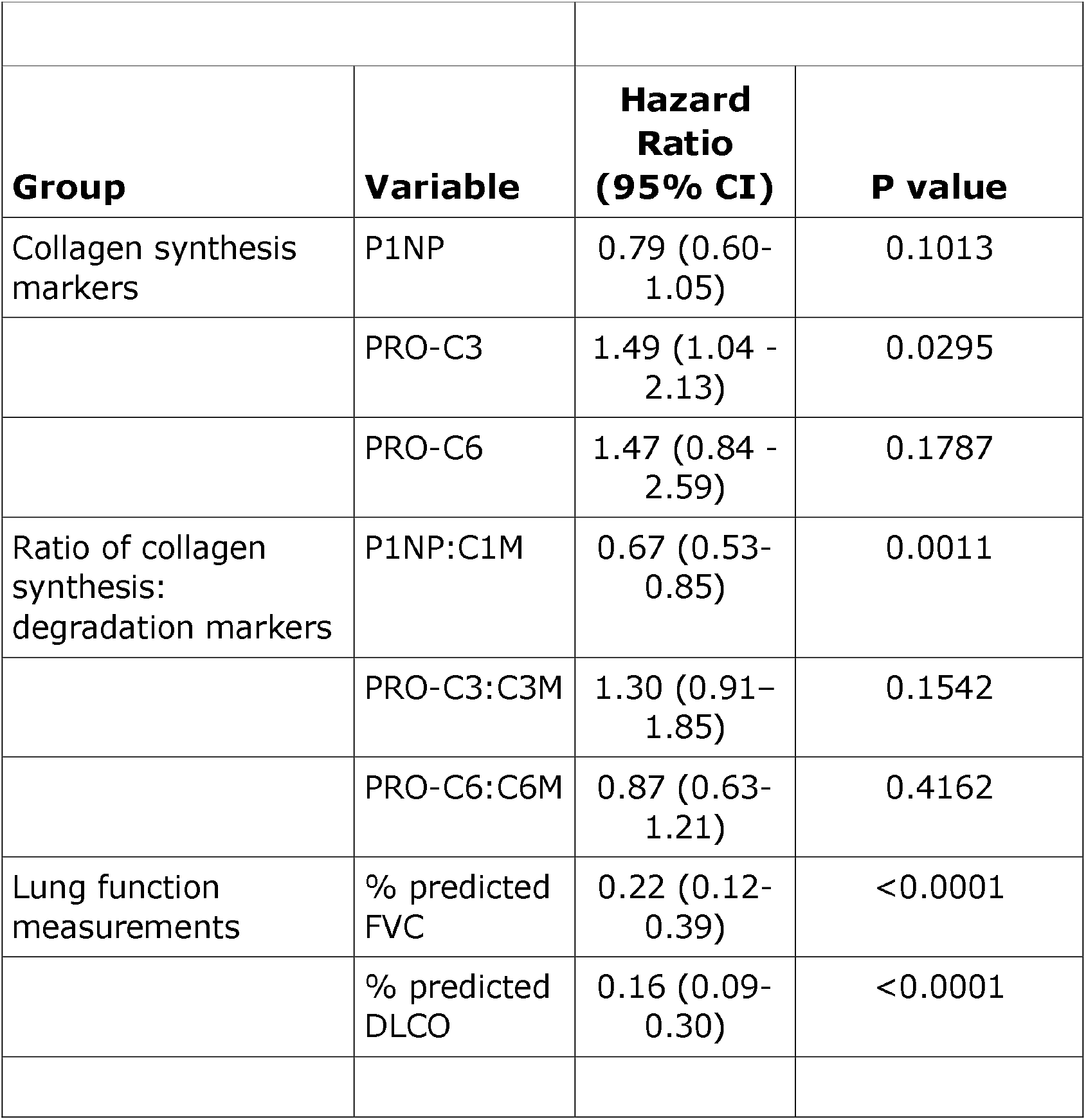
Univariate analysis of overall survival in relation to baseline measures of collagen synthesis neoepitopes for collagen-type 1 (P1NP), - type 3 (PRO-C3) and - type 6 (PRO-C6), the ratio of collagen synthesis: degradation ratios for collagen-type 1 (P1NP:C1M), - type 3 (PRO-C3: C3M), and-type 6 (PRO-C6: C6M) and lung function measurements (% predicted FVC and % predicted DLCO) in IPF subjects. The data are expressed as mean hazard ratio (HR) with 95% CIs for all thresholds and represent the associated change in mortality risk based on a 2-fold increase in the explanatory value.

None of the three synthesis markers were significantly associated with overall survival (**Figure 4**). By contrast, and in keeping with our previously published observations, individuals with rising levels of CRPM, C1M and C6M were at high risk of death compared with those with falling/stable levels (HIGH vs LOW) of CRPM (HR 2.13 [95% CI 1.21 – 3.75] p=0.009, C1M (HR 1.84 [CI 95% 1.03 – 3.27] P=0.038) and C6M (HR 2.19 [1.25 – 3.82], p=0.006) (**Figure 4**). In addition, we also found that rising levels of C3M were associated with higher risk of mortality; (HR 2.44 [CI 95% 1.39 – 4.31] p=0.002) whilst rising levels of the ratio of PRO-C6:C6M was associated with lower mortality (HR 0.55 [0.32-0.95], p=0.032), however, no relationship was observed with levels of P1NP:C1M or PRO-C3:C3M (**Figure 4**).

**Figure 4.**
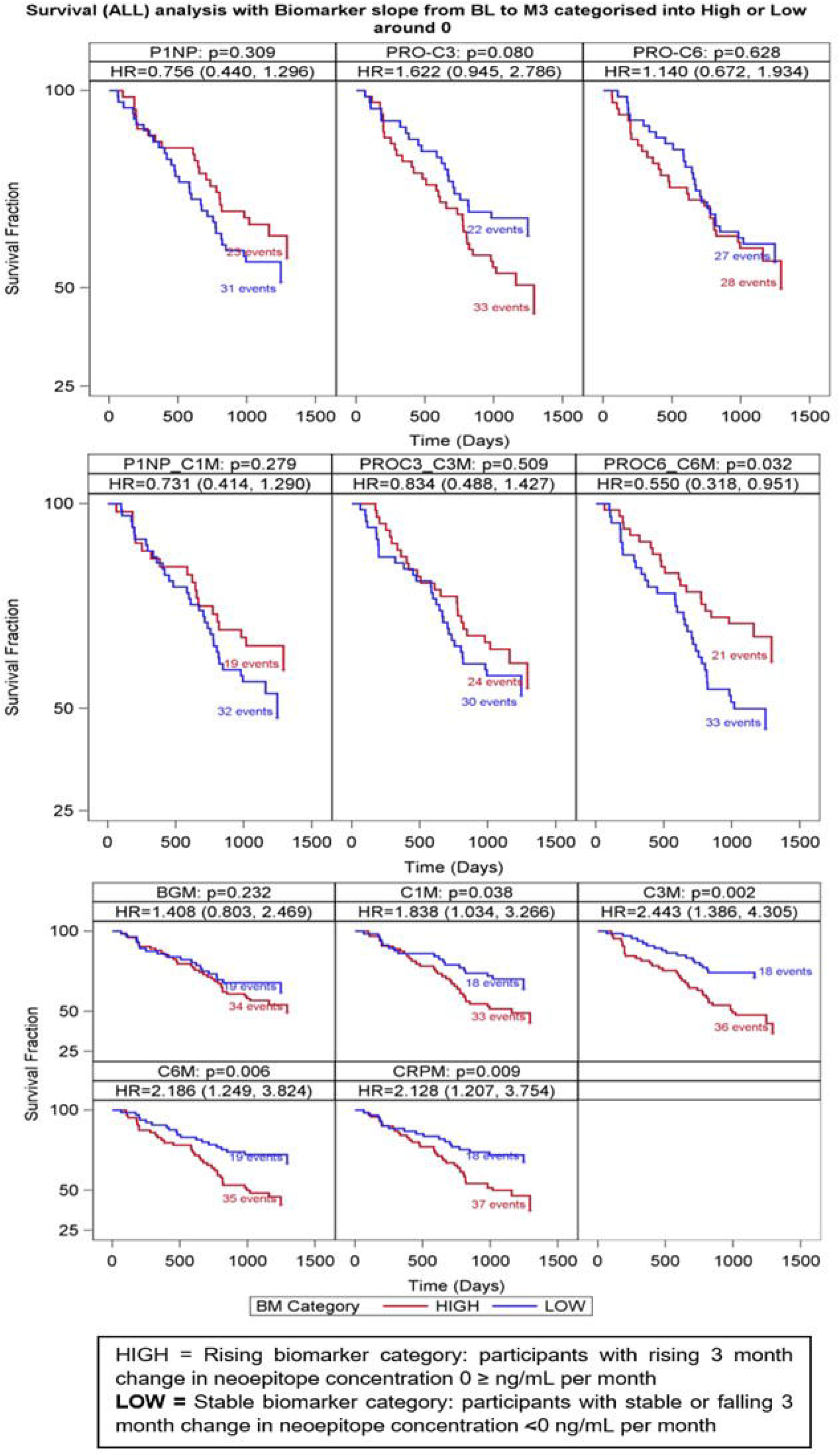
Effect of 3 month change in neoepitope concentrations on overall survival. Subjects with IPF were dichotomised into HIGH or LOW groups, to determine the effect of 3-month change in neoepitope concentration on overall survival, where: HIGH = rising (0>ng/mL per month) neoepitope concentrations from baseline to month 3, >or LOW = stable or falling (<0ng/mL per month) neoepitope concentration from baseline to month 3. For ratio values, the slope values were derived following the creation of the ratio variables and subjects were dichotomised into HIGH or LOW slope groups. Hazard ratio represents the mortality risk in participants with rising neoepitope concentrations relative to those with stable or falling concentrations. Data shown in: top panel - collagen synthesis markers P1NP, PRO-C3, PRO-C6; mid panel - ratio of collagen synthesis/degradation markers: P1NP:C1M, PRO-C3:C3M and PRO-C6:C6M; bottom panel -BGM, C1M, C3M, C6M, CRPM). The number of deaths in each group is shown for each biomarker. P values presented relate to the comparison of the Kaplan-Meir survival curves for each dichotomised measure.

## Discussion

Remodelling of extracellular matrix is pivotal to the development and progression of IPF and results in distinct alterations in collagen turnover. In the current study, we measured synthesis markers of three collagens important in the development of IPF; namely collagen type-1 (P1NP), −3 (PRO-C3) and −6 (PRO-C6) [4, 18, 19]. Given that fibrogenesis in IPF likely reflects an imbalance between synthesis and degradation of collagen, we also measured paired markers of degradation for the same collagens: type-1 (C1M), −3 (C3M) and −6 (C6M). For comparison with our prior observations [7], we also measured degradation of the matrix molecule biglycan (BGM) and C reactive protein (CRPM). Our data demonstrate that synthesis markers for type-3 (PRO-C3) and-6 (PRO-C6) collagen are elevated in subjects with IPF compared with healthy controls and distinguished individuals with progressive disease, suggesting that these biomarkers may reflect underlying pathophysiology driving disease behaviour in IPF.

Collagen type-3, a fibrillar collagen, forms a major component of the lung interstitial extracellular matrix and is synthesised by activated fibroblasts [20]. Our findings support earlier studies demonstrating that synthesis of collagen type-3 is increased in IPF, with high levels of type-3 pro-collagen peptide reported in serum [18] and lavage fluid [21, 22]. In the case of collagen type-3, the synthesis neoepitope, PRO-C3, appears to be a better biomarker of progressive disease than the paired degradation biomarker, C3M. Interestingly, in subjects with more stable disease, the ratio of PRO-C3 to C3M is significantly lower; supporting the idea that synthesis, and increased turnover in general, may be reflective of active remodelling and disease progression.

Alongside collagen type-3, our data suggests that changes in collagen type-6 are also informative of disease behaviour in IPF and may be an important driver of fibrosis. Collagen type-6 filaments are an integral component found at the interface between the basement membrane and interstitial matrix of the lung, and act as an important network for attachment of basement membranes, collagen fibres and cells. Our results support the importance of this collagen in the development of fibrosis. Previous research suggests that collagen type-6 fragments have signalling properties during wound healing and fibrogenesis and may drive poorer outcomes in fibrotic diseases [23]. Compared with other collagens, type-6 collagen has been shown to be a potent inducer of the differentiation and activation of fibroblasts to highly synthetic myofibroblasts; a key pathway leading to the production of collagen and ECM proteins [24]. In addition, increased levels of serum collagen type-6 markers have been demonstrated to act as an early predictor of fibrosis in the liver [15]. This evidence, coupled with data from the current study, suggest that collagen type-6 may in itself be a driver of the fibrotic process, which in turn serves as a useful marker of IPF progression.

Intriguingly, whilst increases in collagen type-1 production are associated with TGFβ-activation of IPF fibroblasts [25], and have been observed in IPF lung [26], we did not find elevated levels of collagen type 1 synthesis neoeptiopes in the serum of IPF subjects. However, collagen type-1 is also the predominant collagen found in bone and it has been estimated that 90% of the synthesis of this collagen measurable in serum is bone derived [27]. This may explain why there was no difference observed in P1NP, the measured marker of collagen type-1 synthesis, between IPF subjects and health controls and why differences were not seen between stable and progressive disease. In contrast to P1NP, the serum degradation marker of collagen type-1, C1M, does not measure bone-derived degradation as the epitope is denatured in bone by osteoclast-released cathepsin K [13]. This disparity in the pools of collagen type-1 being measured may account for the differences observed between this pair of synthesis and degradation neoepitopes.

Consistent with our previous study, baseline levels of C1M, C3M, C6M and CRPM were elevated in subjects with IPF compared with healthy controls. Furthermore, consistent with our prior observations, elevated CRPM distinguished progressive from stable disease from as early as one month after recruitment. In our previous study BGM, C1M, C3M and C6M were also different between individuals with stable and progressive IPF, but these differences only became significant at 6 months. Interestingly, in the current study, the levels of C1M and C6M were significantly higher from 1 month onwards.

Rising levels over 3 months of matrix synthesis markers were not predictive of mortality. This is in distinct contrast with changing levels of the degradation markers CRPM, C1M, C3M and C6M; for each of these markers rising levels over 3 months identified subjects with an increased risk of death. The predictive value of these markers, as shown by the hazard ratios, was consistent with previous results [7]. A changing ratio over 3 months of synthesis and degradation markers for type 6 collagen, PRO-C6:C6M, was also predictive of mortality albeit less strongly than C6M alone. It is possible that the different characteristics of the synthesis and degradation markers may reflect the specific location of individual collagens within the fibrotic lung and their susceptibility to degradation as fibrosis progresses. Alternatively, as is likely to be the case for P1NP, other anatomical pools of collagen turnover, including bone [27] and liver [15], may dilute the serum signal for individual neoepitopes thus limiting their suitability as markers of IPF severity or disease progression.

The absence of reliable and validated short-term measurements that predict disease progression, survival and response to treatment represent a major obstacle in the management of IPF. Physiological measurements, particularly FVC, remain the primary method for monitoring IPF disease progression in both registration clinical trials and clinical practice [28, 29]. Whilst changes in lung function are likely to remain a gold standard for assessing IPF progression, there are several shortcomings of these measurements. Firstly, they are relatively insensitive to subtle changes in disease severity necessitating measurement over longer timeframes (52 weeks in clinical trials) before disease progression can be reliably determined. Although a recent study has demonstrated that daily home spirometry is predictive of disease progression within 3 months, there are some potential limitations with this method, particularly long-term compliance [30]. Secondly, current treatments only slow decline in FVC making determination of treatment response in individuals almost impossible. Serum biomarkers, such as those measured in the current study, have the potential to overcome some of these challenges, by providing a minimally invasive, short-term readout that could supplement FVC measurements to give a more reliable assessment of disease progression and response to anti-fibrotic therapy.

We have shown that neoepitopes reflecting collagen synthesis and degradation in IPF are informative of prognosis, are able to distinguish progressive from non-progressive disease and provide a 3-month read out of change in disease severity. The utility of these observations is currently being tested in a randomised placebo control trial of nintedanib in individuals with IPF and preserved lung function (NCT02788474). If neo-epitopes can be shown to change with therapy, then these markers may be of use in shortening future clinical trials and for determining response to therapy in the clinic.

A major strength of our study is the prospective longitudinal design, which enabled blood sampling to be standardised and enabled collection of robust outcome data. Furthermore, the PROFILE cohort is representative of real-world patients with IPF and includes many subjects who would not have been enrolled in clinical trials. As such our findings are more translatable to clinical practice than insights derived from clinical trial cohorts. The main weakness of this current study is that we undertook analysis of synthesis neoepitopes in a single cohort and did not adhere to a discovery-validation design. However, as this is the only longitudinal study that includes treatment-naive patients with multiple biological samples available that we are aware of, external replication of this study is not currently possible. We feel that this limitation is offset by our demonstration that degradation neo-epitopes behave in a way entirely consistent with our previous observations, which were performed in discovery, and validation sample sets. As such we believe that this confirmation of the usefulness of collagen degradation biomarkers provides indirect validation of our findings and supports our conclusions regarding the potential utility of collagen synthesis biomarkers.

In conclusion, this study demonstrates that collagen type-3 and −6 synthesis biomarkers are elevated in individuals with IPF compared with controls and differ in levels between stable and progressive disease, alongside collagen degradation neoepitopes. In addition, this study re-confirms the value of a panel of ECM degradation biomarkers that are predictive of disease progression and survival in IPF. This panel of biomarkers provides informative insights into matrix turnover and underlying pathophysiology driving disease progression, which may assist in the development of precision approaches to IPF treatment. Furthermore, the predictive prognostic value of change in these biomarkers is discernible within 3 months, providing much needed short-term measurements to assist the identification of at-risk IPF patients, which could be used to supplement physiological assessment to allow for better disease management through targeted and earlier therapeutic intervention strategies.

## Authors’ contributions

RGJ, TMM, RPM, WAF, JKS, EO conceived and designed the study; RB, PLM and participated in recruitment of study patients and collected their data. DJL and MAK developed and undertook the neoepitope assays. EO, AMD, ST, ARK, CBN, LO, WAF, RGJ, and TMM undertook the data analysis. L.O, T.M prepared manuscript. All authors agree with manuscript results and conclusions, made critical revisions and approved final version.

## References

1. Rose D, Montgomery A. Idiopathic pulmonary fibrosis: clinically meaningful primary endpoints in phase 3 clinical trials. Am J Respir Crit Care Med 2013: 187(11): 1269.

2. Hynes RO. The extracellular matrix: not just pretty fibrils. Science 2009: 326(5957): 1216–1219.

3. Selman M, Montano M, Ramos C, Chapela R. Concentration, biosynthesis and degradation of collagen in idiopathic pulmonary fibrosis. Thorax 1986: 41(5): 355–359.

4. Kulkarni T, O’Reilly P, Antony VB, Gaggar A, Thannickal VJ. Matrix Remodeling in Pulmonary Fibrosis and Emphysema. American journal of respiratory cell and molecular biology 2016: Ja.

5. Leeming DJ, Sand JM, Nielsen MJ, Genovese F, Martinez FJ, Hogaboam CM, Han MK, Klickstein LB, Karsdal MA. Serological investigation of the collagen degradation profile of patients with chronic obstructive pulmonary disease or idiopathic pulmonary fibrosis. Biomarker insights 2012: 7: BMI.S9415.

6. McKleroy W, Lee T-H, Atabai K. Always cleave up your mess: targeting collagen degradation to treat tissue fibrosis. American Journal of Physiology-Lung Cellular and Molecular Physiology 2013: 304(11): L709–L721.

7. Jenkins RG, Simpson JK, Saini G, Bentley JH, Russell A-M, Braybrooke R, Molyneaux PL, McKeever TM, Wells AU, Flynn A. Longitudinal change in collagen degradation biomarkers in idiopathic pulmonary fibrosis: an analysis from the prospective, multicentre PROFILE study. The Lancet Respiratory Medicine 2015: 3(6): 462–472.

8. Maher TM, Oballa E, Simpson JK, Porte J, Habgood A, Fahy WA, Flynn A, Molyneaux PL, Braybrooke R, Divyateja H. An epithelial biomarker signature for idiopathic pulmonary fibrosis: an analysis from the multicentre PROFILE cohort study. The Lancet Respiratory Medicine 2017: 5(12): 946–955.

9. Raghu G, Collard HR, Egan JJ, Martinez FJ, Behr J, Brown KK, Colby TV, Cordier J-FF, Flaherty KR, Lasky JA, Lynch DA, Ryu JH, Swigris JJ, Wells AU, Ancochea J, Bouros D, Carvalho C, Costabel U, Ebina M, Hansell DM, Johkoh T, Kim DS, King TE, Kondoh Y, Myers J, Müller NL, Nicholson AG, Richeldi L, Selman M, Dudden RF, Griss BS, Protzko SL, Schünemann HJ, Fibrosis AEJACoIP. An official ATS/ERS/JRS/ALAT statement: idiopathic pulmonary fibrosis: evidence-based guidelines for diagnosis and management. American journal of respiratory and critical care medicine 2011: 183(6): 788–824.

10. Leeming DJ, Larsen D, Zhang C, Hi Y, Veidal S, Nielsen R, Henriksen K, Zheng Q, Barkholt V, Riis B. Enzyme-linked immunosorbent serum assays (ELISAs) for rat and human N-terminal pro-peptide of collagen type I (P1NP)—assessment of corresponding epitopes. Clinical biochemistry 2010: 43(15): 1249–1256.

11. Nielsen MJ, Nedergaard AF, Sun S, Veidal SS, Larsen L, Zheng Q, Suetta C, Henriksen K, Christiansen C, Karsdal MA. The neo-epitope specific PRO-C3 ELISA measures true formation of type III collagen associated with liver and muscle parameters. American journal of translational research 2013: 5(3): 303.

12. Nedergaard A, Sun S, Karsdal MA, Henriksen K, Kjær M, Lou Y, He Y, Zheng Q, Suetta C. Type VI collagen turnover¤related peptides—novel serological biomarkers of muscle mass and anabolic response to loading in young men. Journal of cachexia, sarcopenia and muscle 2013: 4(4): 267–275.

13. Leeming DJ, He Y, Veidal S, Nguyen Q, Larsen D, Koizumi M, Segovia-Silvestre T, Zhang C, Zheng Q, Sun S. A novel marker for assessment of liver matrix remodeling: anenzyme-linked immunosorbent assay (ELISA) detecting a MMP generated type I collagen neo-epitope (C1M). Biomarkers 2011: 16(7): 616–628.

14. Barascuk N, Veidal S, Larsen L, Larsen D, Larsen M, Wang J, Zheng Q, Xing R, Cao Y, Rasmussen LM. A novel assay for extracellular matrix remodeling associated with liver fibrosis: An enzyme-linked immunosorbent assay (ELISA) for a MMP-9 proteolytically revealed neo-epitope of type III collagen. Clinical biochemistry 2010: 43(10-11): 899–904.

15. Veidal SS, Karsdal MA, Vassiliadis E, Nawrocki A, Larsen MR, Nguyen QHT, Hägglund P, Luo Y, Zheng Q, Vainer B. MMP mediated degradation of type VI collagen is highly associated with liver fibrosis-identification and validation of a novel biochemical marker assay. PLoS One 2011: 6(9): e24753.

16. Skjøt-Arkil H, Schett G, Zhang C, Larsen DV, Wang Y, Zheng Q, Larsen MR, Nawrocki A, Bay-Jensen A-C, Henriksen K. Investigation of two novel biochemical markers of inflammation, matrix metalloproteinase and cathepsin generated fragments of C-reactive protein, in patients with ankylosing spondylitis. Clinical and experimental rheumatology 2012: 30(3): 371–379.

17. Cotes J, Chinn D, Quanjer PH, Roca J, Yernault J-C. Standardization of the measurement of transfer factor (diffusing capacity). Eur Respiratory Soc, 1993.

18. Kirk J, Bateman E, Haslam P, Laurent GJ, Turner-Warwick M. Serum type III procollagen peptide concentration in cryptogenic fibrosing alveolitis and its clinical relevance. Thorax 1984: 39(10): 726–732.

19. Specks U, Nerlich A, Colby TV, Wiest I, Timpl R. Increased expression of type VI collagen in lung fibrosis. American journal of respiratory and critical care medicine 1995: 151(6): 1956–1964.

20. Frantz C, Stewart KM, Weaver VM. The extracellular matrix at a glance. J Cell Sci 2010: 123(24): 4195–4200.

21. Low RB, Giancola MS, King T, Chapitis J, Vacek P, Davis GS. Serum and bronchoalveolar lavage n-terminal type III procollagen peptides in idiopathic pulmonary fibrosis. American Review of Respiratory Disease 1992: 146: 701–701.

22. Bjermer L, Lundgren R, Hällgren R. Hyaluronan and type III procollagen peptide concentrations in bronchoalveolar lavage fluid in idiopathic pulmonary fibrosis. Thorax 1989: 44(2): 126–131.

23. Karsdal M, Nielsen S, Leeming D, Langholm L, Nielsen M, Manon-Jensen T, Siebuhr A, Gudmann N, Rønnow S, Sand J. The good and the bad collagens of fibrosis-Their role in signaling and organ function. Advanced drug delivery reviews 2017: 121: 43–56.

24. Naugle JE, Olson ER, Zhang X, Mase SE, Pilati CF, Maron MB, Folkesson HG, Horne WI, Doane KJ, Meszaros JG. Type VI collagen induces cardiac myofibroblast differentiation: implications for postinfarction remodeling. American Journal of Physiology-Heart and Circulatory Physiology 2006: 290(1): H323–H330.

25. Ramos C, Moñtano M, Garciá-Alvarez J. Fibroblasts from idiopathic pulmonary fibrosis and normal lungs differ in growth rate, apoptosis, and tissue inhibitor of metalloproteinases expression. American journal of … 2001.

26. Raghu G, Striker LJ, Hudson LD, Striker GE. Extracellular matrix in normal and fibrotic human lungs. Am Rev Respir Dis 1985: 131(2): 281–289.

27. Leeming D, Alexandersen P, Karsdal M, Qvist P, Schaller S, Tanko L. An update on biomarkers of bone turnover and their utility in biomedical research and clinical practice. European journal of clinical pharmacology 2006: 62(10): 781–792.

28. Lee SH, Shim HS, Cho SH, Kim SY, Lee SK, Son JY, Jung JY, Kim EY, Lim JE, Lee KJ, Park BH, Kang YA, Kim YS, Kim SK, Chang J, Park MS. Prognostic factors for idiopathic pulmonary fibrosis: clinical, physiologic, pathologic, and molecular aspects. Sarcoidosis Vasc Diffuse Lung Dis 2011: 28(2): 102–112.

29. du Bois R, Nathan S, Richeldi L, Schwarz M, Noble P. Idiopathic pulmonary fibrosis: lung function is a clinically meaningful endpoint for phase III trials. Am J Respir Crit Care Med 2012: 186(8): 712–715.

30. Russell A-M, Adamali H, Molyneaux PL, Lukey PT, Marshall RP, Renzoni EA, Wells AU, Maher TM. Daily home spirometry: an effective tool for detecting progression in idiopathic pulmonary fibrosis. American journal of respiratory and critical care medicine 2016: 194(8): 989–997.

