## Supplementary figures for "Biomarkers of collagen synthesis predict progression in the PROFILE idiopathic pulmonary fibrosis cohort"

**Online Data Supplement**

| Assay name | Target | Antibody type | Detection range, LLOQ-ULOQ (ng/mL) | Intra-assay variation (%) | Inter-assay variation (%) | Assay principle reference |
| --- | --- | --- | --- | --- | --- | --- |
| C1M | MMP-2/9/13 degraded type I collagen | Monoclonal | 10-200 | 2.7-8.2 | 5.5-18.0 | (12) |
| C3M | MMP-9 degraded type III collagen | Monoclonal | 4-88 | 6.6-15.1 | 2.0-4.1 | (13) |
| C6M | MMP-2/9 degraded type vI collagen | Monoclonal | 6-267 | 2.0-8.0 | 4.0-18.0 | (32) |
| CRPM | MMP-1/9 degraded C-reactive protein | Monoclonal | 2.0-72.0 | 2.2-6.0 | 4.1-21.1 | (14) |
| BGM | MMP-9 degraded biglycan | Monoclonal | 4.1-190.0 | 2.0-6.0 | 5.0-20.0 | (33) |
| P1NP | N-terminal propeptide of type I collagen (formation marker) | Monoclonal | 14.0-516.0 | 1.0-8.0 | 3.0-13.0 | (9) |
| PRO-C3 | N-terminal propeptide of type III collagen (formation marker) | Monoclonal | 2.6-116.0 | 1.8-9.3 | 8.0-12.0 | (10) |
| PRO-C6 | C-terminal of type VI collagen (formation marker) | Monoclonal | 0-8-134.0 | 1.1-5.3 | 3.4-12.4 | (34) |

**Table 1E. Detailed outline of the target and assay performance characteristics for the ELISA assays specific to each neoepitope. The upper and lower limits of quantification, as well as intra- and inter-assay variation is also defined for each neoepitope specific assay. LLOQ= lower limit of quantification, ULOQ= Upper limit of quantification.**

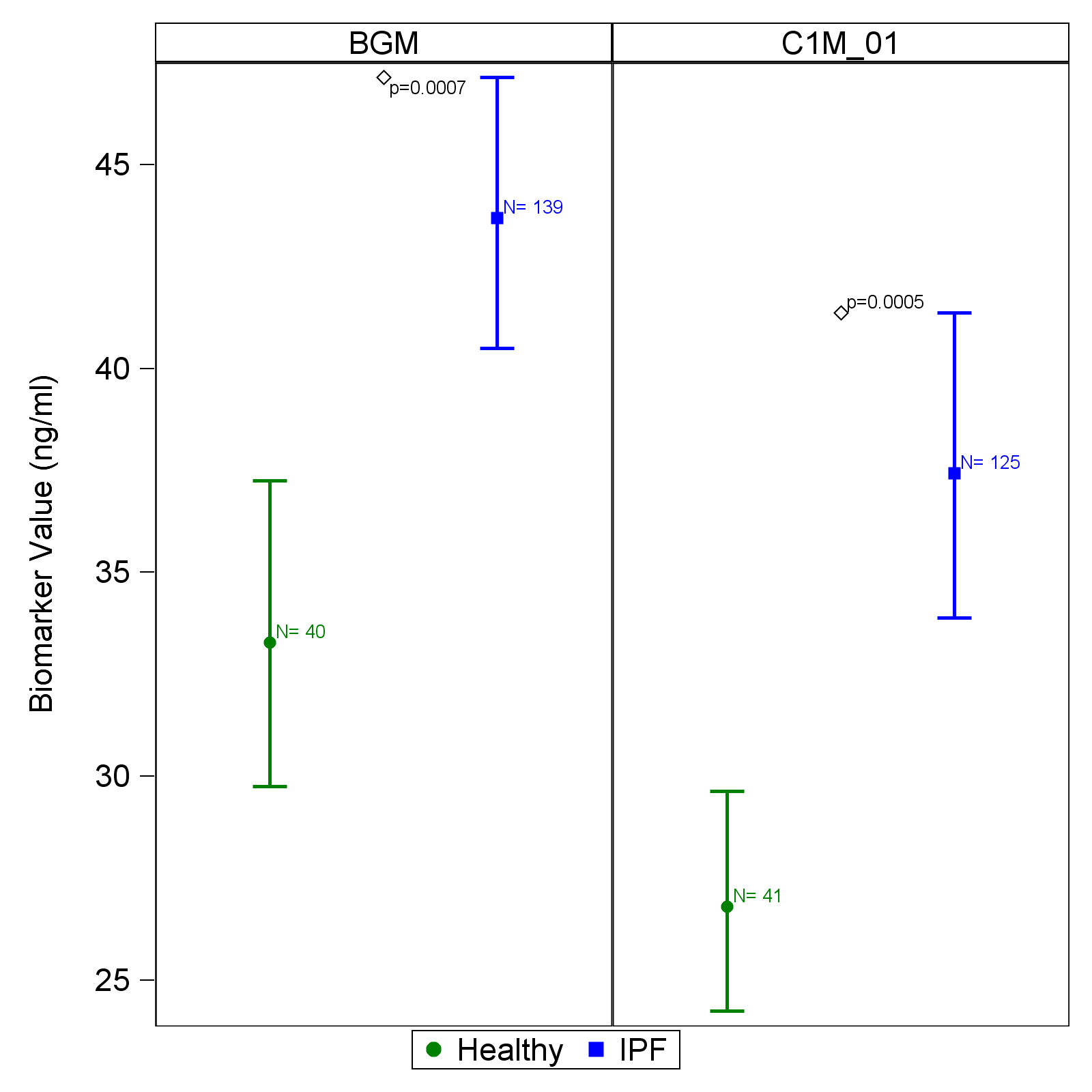

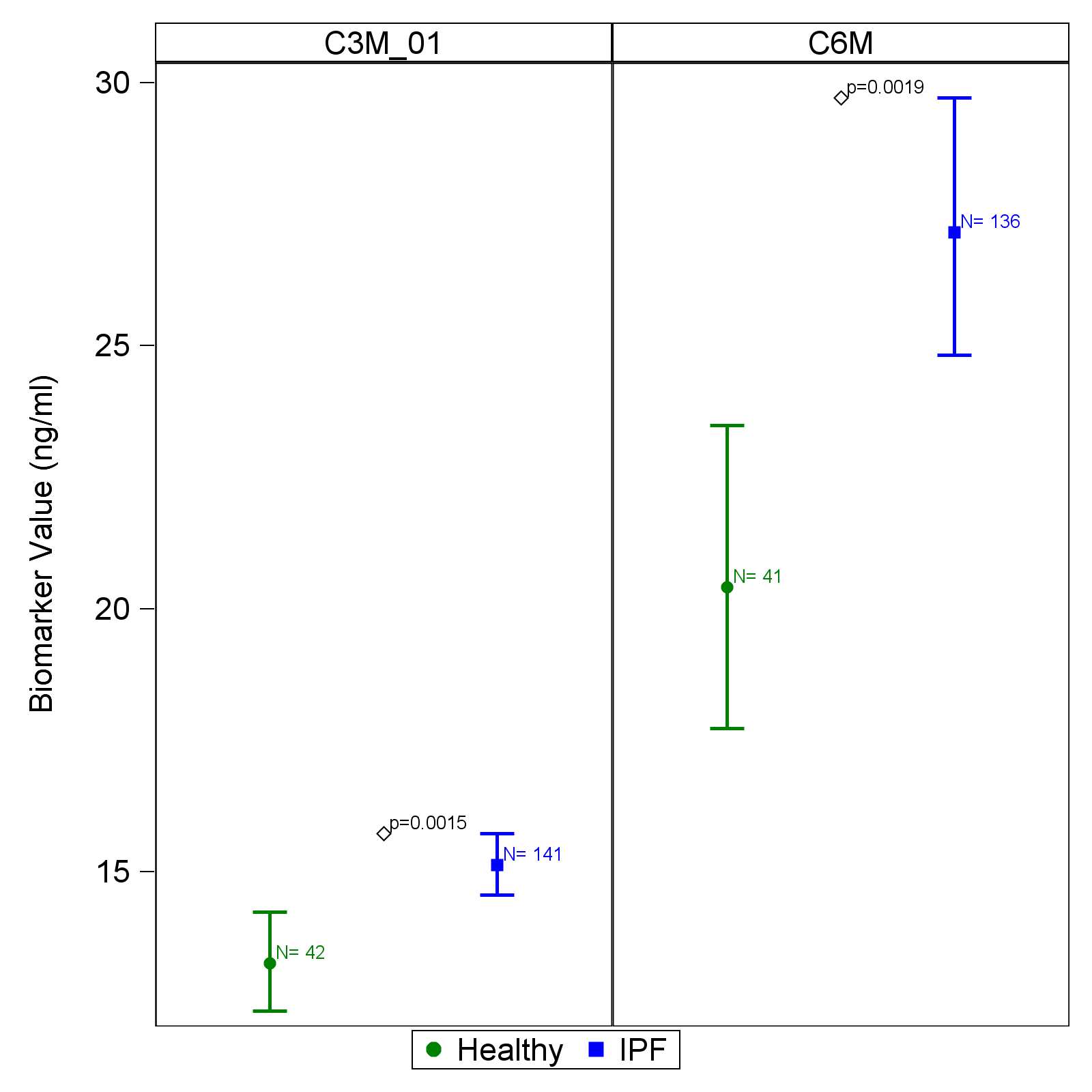

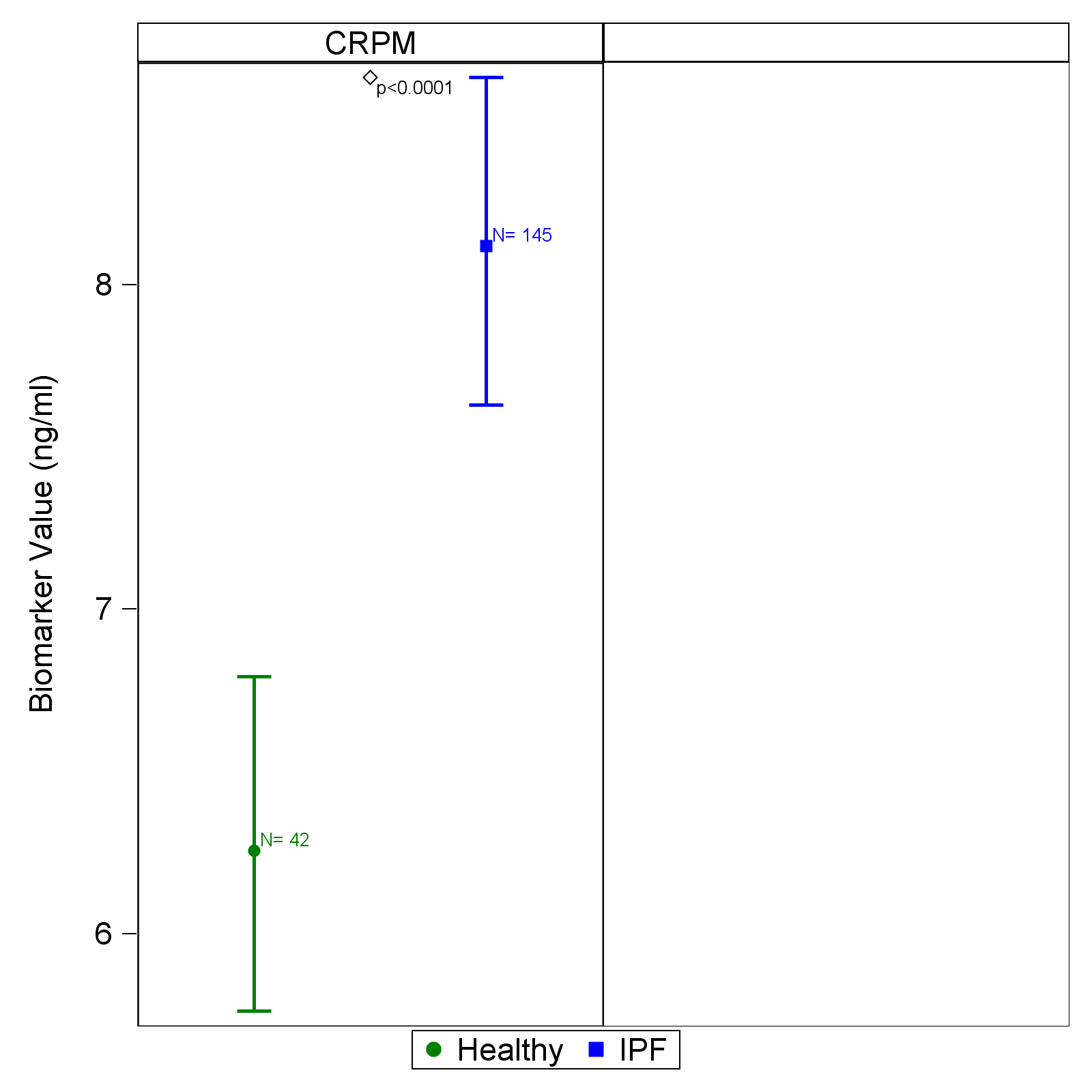

**Figure 1E. Baseline comparison of collagen degradation neoepitope (BGM, C1M, C3M, C6M and CRPM) concentrations in healthy controls (n=50) and participants with idiopathic pulmonary fibrosis (n=145).**

Plots represent mean and 95% CI (error bars) adjusted for gender.

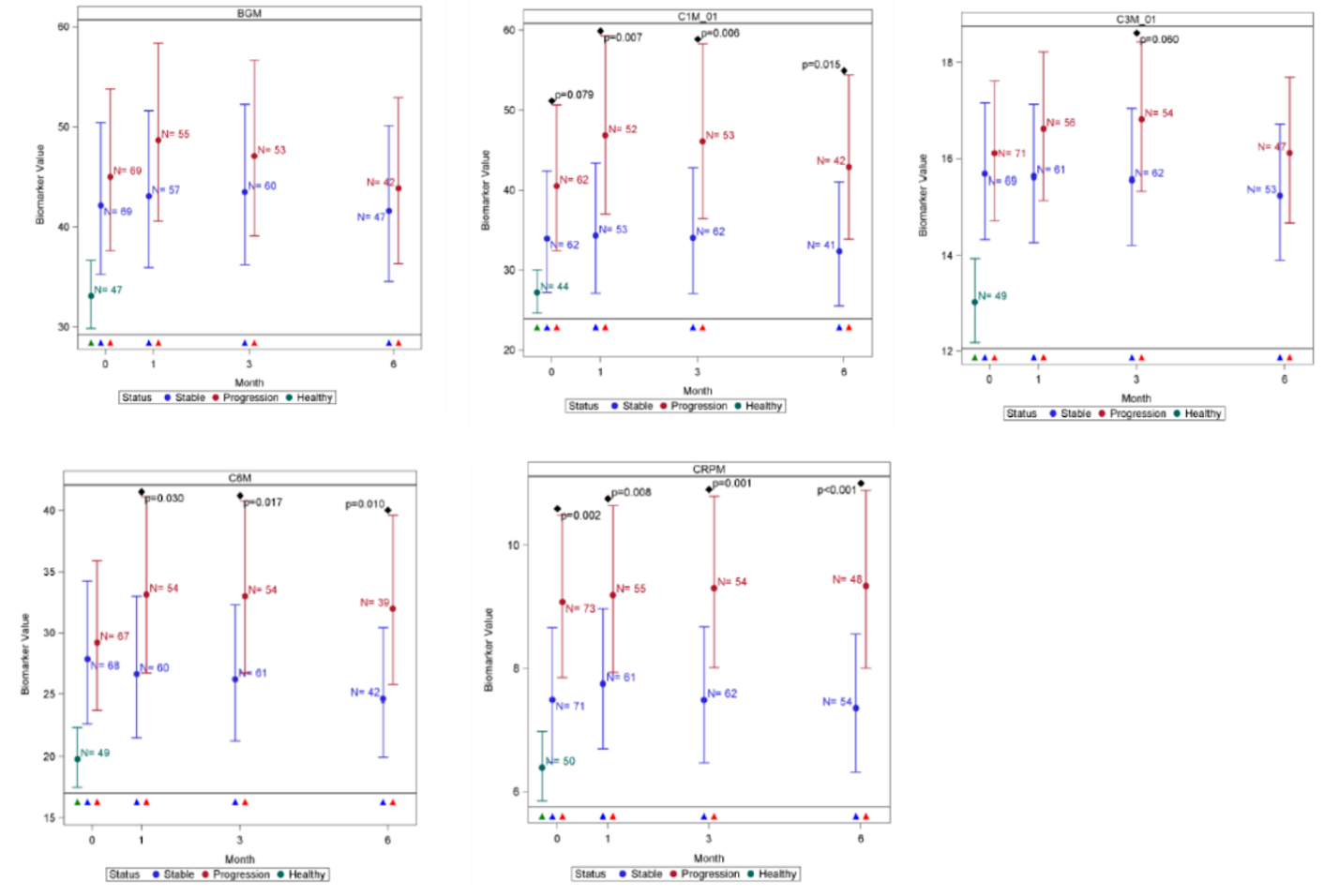

**Figure 2E. Comparison of neoepitope concentrations in healthy controls (green) and participants with stable (blue) and progressive (red) idiopathic pulmonary fibrosis at baseline and subsequently at 1-, 3- and 6-months.**

Plots represent mean and 95% CI (error bars) adjusted for age, sex, site and smoking status. Disease progression was defined as all-cause mortality or ≧10% decline in forced vital capacity at 12 months. The number of evaluable samples available for analysis at each time point are provided in the graph. P values are provided where significant (p<0.05) differences were observed between stable and progressive disease at a particular time point.
